# MUUMI: an R package for statistical and network-based meta-analysis for MUlti-omics data Integration

**DOI:** 10.1101/2025.03.10.642416

**Authors:** Simo Inkala, Michele Fratello, Giusy del Giudice, Giorgia Migliaccio, Angela Serra, Dario Greco, Antonio Federico

## Abstract

Disentangling physiopathological mechanisms of biological systems through high-level integration of omics data has become a standard procedure in life sciences. However, platform heterogeneity, batch effects, and the lack of unified methods for single- and multi-omics analyses represent relevant drawbacks that hinder the extrapolation of a meaningful biological interpretation. Statistical meta-analysis is widely used in order to integrate several omics datasets of the same type, leading to the extrapolation of robust molecular signatures within the investigated system. Conversely, statistical meta-analysis does not allow the simultaneous investigation of different molecular layers, and, therefore, the integration of multi-modal data deriving from multi-omics experiments. Although in the last few years a number of valid tools designed for multi-omics data integration have emerged, they have never been combined with statistical meta-analysis tools in a unique analytical solution in order to support meaningful biological interpretation. Network science is at the forefront of systems biology, where the inference of molecular interactomes allowed the investigation of perturbed biological systems, by shedding light on the disrupted relationships that keep the homeostasis of complex systems. Here, we present MUUMI, an R package that unifies network-based data integration and statistical meta-analysis within a single analytical framework. MUUMI allows the identification of robust molecular signatures through multiple meta-analytic methods, inference and analysis of molecular interactomes and the integration of multiple omics layers through similarity network fusion. We demonstrate the functionalities of MUUMI by presenting two case studies in which we analysed 1) 17 transcriptomic datasets on idiopathic pulmonary fibrosis (IPF) from both microarray and RNA-Seq platforms and 2) multi-omics data of THP-1 macrophages exposed to different polarising stimuli. In both examples, MUUMI revealed biologically coherent signatures, underscoring its value in elucidating complex biological processes.

**Availability and implementation:** MUUMI is freely available at https://github.com/fhaive/muumi.

## Background

Omics data integration has long posed substantial challenges, including substantial variability in data quality, lack of standardization, and the inherent heterogeneity of platforms and data formats. These issues can introduce noise, bias, and limited interoperability, complicating efforts to combine results across multiple studies. Remarkable efforts have been carried out in order to make omics data more integrable (1, 2, 3, 4, 5). Nevertheless, methods for interpreting integrated omics data remain scattered, and the research community still lacks a comprehensive resource that unifies both data integration and downstream interpretation.

Statistical meta-analysis has for long represented a crucial method to extrapolate evidence through the integration of independent studies addressing the same research question, aiming to generate a quantitative estimate of the phenomenon under consideration (6). Meta-analysis can be performed at the aggregate data (AD) level—pooling summary statistics for each study—or at the individual participant data (IPD) level—integrating raw data for each participant (7). While IPD meta-analyses are often considered ideal, practical limitations such as data availability, intellectual property constraints, and high costs continue to make AD-based meta-analysis the most widely adopted approach (8). Over the years, several software solutions have been developed for clinical meta-analyses including the Excel plugins MetaXL (Barendregt and Doi, 2009) and Mix 2.0 (Bax, 2016), Revman (Cochrane Collaboration, 2011), Comprehensive Meta-Analysis Software (CMA (Borenstein et al., 2005)), JASP (JASP Team, 2018) and MetaFOR library for R (Viechtbauer, 2010). While these packages can be adapted to basic science projects, difficulties may arise due to large and complex datasets and heterogeneity in experimental methodology (6). Therefore, a pressing need for tools specifically tailored to multi-study omics integration still remains.

Network science provides a complementary and increasingly popular framework for deciphering the intricate relationships among genes, proteins, or metabolites in complex biological systems (9, 10, 11). The inference of these molecular interactomes finds diverse applications, including biomarker discovery, patient stratification, and druggability assessment. By shifting the focus from individual molecular entities to broader connectivity patterns, network-based approaches can reveal functional modules, regulatory relationships, and disease-associated sub-networks that may be overlooked by traditional methods. While network modelling has been widely utilised to integrate multi-view biological datasets (1), its potential in supporting and complementing traditional meta-analytical approaches has not yet fully exploited.

In this study, we developed MUUMI, an R package designed to streamline omics data integration and interpretation through both statistical and network-based meta-analysis. MUUMI combines multiple meta-analysis techniques within a unified pipeline, complemented by tools for network inference, community detection, and functional annotation. Moreover, it supports multi-omics data integration through Similarity Network Fusion (1), extrapolating molecular signals across distinct omics layers. We showcase the functionalities of MUUMI in two case studies: 1) a single-omics, multi-study integrative analysis of idiopathic pulmonary fibrosis (IPF) datasets, and 2) a multi-omics dataset investigating macrophage polarization. Figure 1 shows the functionalities of the MUUMI package.

**Figure 1.**
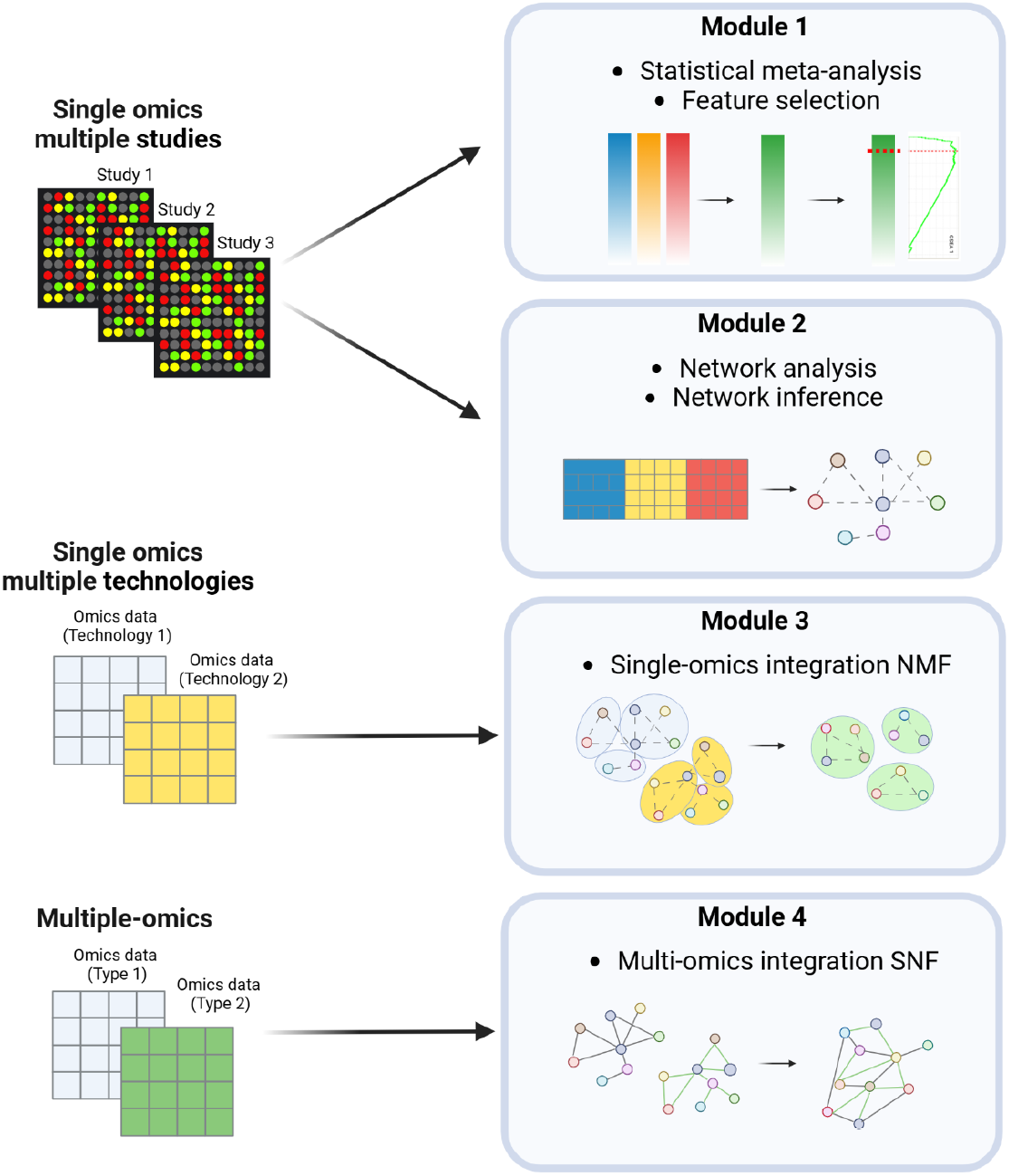
Overview of the functionalities of the MUUMI package.

## Implementation

The MUUMI package comprises four main modules. Module 1 focuses on statistical meta-analysis, allowing the user to pool differential analysis results from multiple independent studies addressing the same biological question. It produces a ranked list of molecular entities (e.g., genes, proteins, etc.) by integrating three distinct meta-analytic approaches: effect size, p-value-based, and rank product. The effect size method accounts for both within- and between-study variability (12), while Fisher’s sum of logs method aggregates p-values from multiple independent studies (13). Rank product, on the other hand, is a non-parametric statistical approach that merges differential analysis outcomes from individual studies based on within-study ranking of molecular entities; these ranks are then combined via a one-class analysis of the rank-product statistic (14, 15). Beyond these individual methods, MUUMI provides the opportunity to run an ensemble of these methods to increase the robustness, comprehensiveness, statistical power, and reproducibility of the analysis (16). Module 1 also includes a feature selection function that derives a threshold for the meta-analysis rank based on the biological significance of molecular entities.

This is accomplished through a Gene Set Enrichment Analysis (GSEA)-based approach, which calculates enrichment scores of ranked entities over pathway-specific gene sets. The resulting threshold provides a biologically informed cutoff for selecting features for downstream analyses.

Module 2 is dedicated to network analysis and inference. Since omics datasets commonly originate from multiple sources and experimental platforms, batch effects can hide genuine biological signals. MUUMI addresses this challenge with a function that aggregates, scales, and mitigates batch effects, producing an integrated matrix for use in further analyses, such as network inference. Subsequently, users can infer robust biological networks using an ensemble of methods (17). Various analyses can then be performed on these inferred networks, including community detection via walktrap (18), louvain (19), spinglass (20, 21), or the greedy algorithm (22) as well as module-specific functional annotation and visualization.

Module 3 facilitates single-omics data integration derived from diverse platforms and technologies (e.g., RNA-Seq and microarray data). This module enables the aggregation of networks obtained from different single-omics views. By leveraging non-negative matrix factorization (NMF), MUUMI generates a unified set of community labels from separately constructed molecular networks, aiding in the cross-platform comparison and interpretation of network structures.

Finally, Module 4 extends the integration paradigm to a multi-omics setting through the Similarity Network Fusion (SNF) algorithm (1). SNF constructs a similarity network for each omics layer (e.g., transcriptomics, proteomics, epigenomics) and iteratively fuses these into a single consensus network. This unified representation highlights relationships consistent across multiple data types.

To further assist in the interpretation and visualization of constructed networks, this final module also supports overrepresentation analysis (ORA) on network communities and a topological enrichment approach based on Edge Set Enrichment Analysis (ESEA). Results can be visualized at both the individual and comparative levels for single or multiple networks, facilitating a deeper understanding of complex molecular interactions and enriched functional pathways.

## Application

We illustrate the functionalities of MUUMI through two distinct case studies. In the first, we analyzed and integrated 17 transcriptomics datasets derived from both RNA-Seq and DNA microarray platforms. The datasets, publicly available on Zenodo (https://zenodo.org/doi/10.5281/zenodo.10692128), encompass lung biopsies from idiopathic pulmonary fibrosis (IPF) patients and healthy controls. Our primary objective was to identify robust IPF-associated genes using a statistical meta-analysis across all datasets. This analysis ranked a set of genes, notably including several collagens, consistent with the pathological extracellular matrix (ECM) deposition, typical of lung fibrosis. The meta-analysis also highlighted several metalloproteinases, whose dysregulation leads to significant tissue remodeling within the lung microenvironment.

Building on these findings, we separately aggregated gene expression measurements from the RNA-Seq and DNA microarray datasets to construct co-expression networks for both IPF and healthy samples. By comparing the networks derived exclusively from RNA-Seq data with those obtained through integration of RNA-Seq and microarray data, we observed enhanced functional characterization in the integrated network. Specifically, the integrated network modules more readily pinpointed processes central to IPF, including ECM degradation, organelle biogenesis and maintenance, and cell junction organization.

In the second case study, we examined macrophage plasticity by integrating transcriptomics and DNA methylation data following exposure to polarizing stimuli (IL4-IL13 and LPS-IFNγ). This multi-omics approach revealed mechanistic signatures underlying both short- and long-term molecular remodeling. Under LPS-IFNγ stimulation, we observed strong correlations among genes involved in TLR and TNF signaling pathways, consistent with known pro-inflammatory cascades (23). Specifically, LPS binds TLR4, triggering MyD88-dependent signaling that activates NF-κB and induces TNF-α expression, thereby amplifying inflammation through autocrine and paracrine mechanisms. Conversely, the IL4-IL13 network exhibited fewer enriched pathways and remained more similar to unstimulated macrophages, indicating a comparatively milder response to IL4-IL13 than to LPS-IFNγ.

## Discussion

Despite many methodologies for multi-omics data integration and meta-analysis have been developed, translating these efforts into biologically meaningful, mechanistic insights remains challenging. Existing softwares focus on isolated aspects of meta-analysis, disregarding the power of network-based approaches to give an holistic view of the biological system or condition under investigation. Other tools provide complex network-based methodologies, but lack functions to aid a robust interpretation of the results. These limitations collectively hinder the exploitation of omics data integration to generate actionable biological insights.

We developed MUUMI to address this gap, by integrating multiple meta-analytical approaches with a comprehensive suite of network-based functionalities, spanning from the inference to the analysis and interpretation of the networks. Moreover, the capacity of MUUMI to handle both single-omics and multi-omics datasets facilitates a more complete interpretation of complex biological systems, providing deeper insights into disease mechanisms, gene regulation, or biomarker discovery. MUUMI includes several exposed functions that can be combined in different ways so as to allow a full customisation of the analysis pipelines according to the biological condition under investigation. In addition, the network-driven interpretation steps, including module functional annotation and the topological enrichment analyses, provide valuable tools to evaluate and explore the biological relevance of the results. The two case studies presented—an integrative analysis of IPF datasets across RNA-Seq and microarray platforms, and a multi-omics study of macrophage plasticity upon polarising stimuli—demonstrate how MUUMI can reveal biologically plausible insights while accommodating heterogeneous data sources.

## Conclusion

We developed MUUMI, an R package leveraging omics data meta-analysis, integration and interpretation that implements traditional and network-based approaches to unleash the power of multi-study datasets. Statistical and network-based approaches are integrated in a unique framework, allowing the user to derive robust and biologically meaningful results from different studies. The applicability and the functionalities of MUUMI were showcased in the performed case studies.

## Supporting information

MUUMI implementation

MUUMI case studies

