## Supplementary material for "MUUMI: an R package for statistical and network-based meta-analysis for MUlti-omics data Integration": MUUMI implementation

MUUMI is an R package organised in 4 modules. This package is suitable for researchers looking for an easy solution to integrate single omics multi-study datasets and carry out both statistical and network-based meta-analysis. Following are reported Bioconductor and CRAN R packages used in MUUMI:

clusterProfiler >= 3.14.3
doParallel >= 1.0.16
dplyr >= 1.0.2
esc >= 0.5.1 ESEA >= 1.0
fgsea >= 1.12.0
foreach >= 1.4.4
ggplot2 >= 3.3.5
gplots >= 3.0.1.1
grid >= 3.6.3
gridExtra >= 2.3
igraph >= 1.2.6
matrixStats >= 0.59.0
metafor >= 3.0-2
metap >= 1.5
minet >= 3.44.1
nmfbin >= 0.2.1
parallel >= 3.6.3
plyr >= 1.8.4
RankProd >= 3.12.0
ReactomePA >= 1.30.0
reshape2 >= 1.4.3
SNFtool >= 2.3.1
stringr >= 1.4.0
tidyr >= 1.1.2
TopKLists >= 1.0.7

**Module 1 - Meta-analysis of multi-study omics datasets.**

In module 1, the user can perform a meta-analysis with the following methods: effect size (1), sum of logs (Fisher's method) and rank product (2,3). The functions to run the meta-analysis with such methods are *calc_effect_size_rank*, *calc_pvalue_based_rank* and *calc_rank_base_rank*, respectively. MUUMI also encompasses a function to perform a meta-analysis with an ensemble of multiple methods named *run_ensembl_metanalysis*.
The input required for all the functions to perform the statistical meta-analysis is a data frame reporting genes on the rows and adjusted p-values for each dataset on the columns. The p-values originate from differential analysis between the biological condition of interest and the control. One example could be the case of RNASeq datasets where, upon differential expression analysis (carried out with tools such as DESeq2 (4), EdgeR (5), NOISeq (6), etc.) the user obtains adjusted p-values for all the genes reported in the annotation. Similarly, for DNA microarray analysis, the p-values could derive from the output of a differential analysis performed through the use of limma (7). The second input of the functions is a vector called "class" that includes the class labels for each sample. In the case of a two-class unpaired design, the class label can be either 0 (representing the control group) or 1 (representing the case group). For one-class data, the label for each sample should be 1. The third input is a vector called "origin" that contains the origin labels for each sample (i.e. the dataset ID).
The *calc_effect_size_rank* function takes into account the mean adjusted p-value, sample sizes (as a vector representing the number of genes in each sample), and gene names to calculate the effect size for each gene by using the chi-square method (according to the *effect_sizes* function from the esc package). Effect size measures the magnitude of differences, providing an estimate of the practical significance of the difference. Effect size is a standardized measure, making it easier to compare across different studies and data sets (8,9). However, effect size can be influenced by sample size, so it can be sensitive to outliers or small numbers of replicates. Besides, effect size does not provide information about the statistical significance of the difference (10, 11).
By using the function *calc_pvalue_based_rank*, a p-value based rank is calculated by aggregating the adjusted p-values for each gene using the Fisher's sum of logs method. This function implements the *sumlog* function from the metap package (Dewey M (2022). *metap: meta-analysis of significance values*. R package version 1.8.). While p-values provide a universal measure of statistical significance, they are severely affected by the sample size, being sensitive to outliers or small numbers of replicates. The function *calc_rank_base_rank* computes the rank product statistic on the p-values for each gene across all datasets by giving in output a rank product-based gene rank. The function implements the *RP.advance* function from the RankProd package (3,12). The output of *RP.advance* is used to internally compute

the final gene rank through the use of the *Borda* function from the TopKLists package (https://doi.org/10.1515/sagmb-2014-0093). Rank product is a robust non-parametric method that can handle violations of assumptions about normality or equal variance. Moreover, rank product does not require a pre-determined threshold for statistical significance, making it appropriate for meta-analyses with diverse datasets. This method is based on biological reasoning related to the fold-change (FC) criterion. It identifies genes that consistently exhibit the strongest upregulation or downregulation across several replicate experiments. Additionally, it provides a solution to overcome the differences among multiple datasets, making it applicable for meta-analysis. One disadvantage of the rank product ranking is that it assumes that the effects being measured are additive, so it may not be appropriate for data with interactions or non-linear relationships.
In addition, the MUUMI package offers a function that implements all of the three above-mentioned meta-analysis methods named *run_ensembl_metanalysis*. In addition to the input file (as explained above for the previous functions), the user needs to provide 1) one or more of the following flags in order to choose the method(s) to include in the ensemble analysis: "*effect_size*", "*pvalue*", "*rank_product*", and 2) the metric to use in order to compute the aggregated gene rank (“*mean*”, “*median*”, “*geometric mean*”, and “*L2 norm*”, according to the Borda method). The meta-analysis pipeline is illustrated in Figure S1.
Integrating rank information from the three methods by performing the ensemble meta-analysis offers several benefits. Using multiple methods can increase the statistical power of the analysis, providing robustness to the results. Moreover, by utilizing this approach, it is possible to identify genes that are genuinely associated with the phenotype of interest, increasing the confidence of their biological relevance and the reproducibility of the results. Overall, using an ensemble of methods to compute gene ranks from multiple studies can increase the robustness, comprehensiveness, statistical power, and reproducibility of the analysis (13).

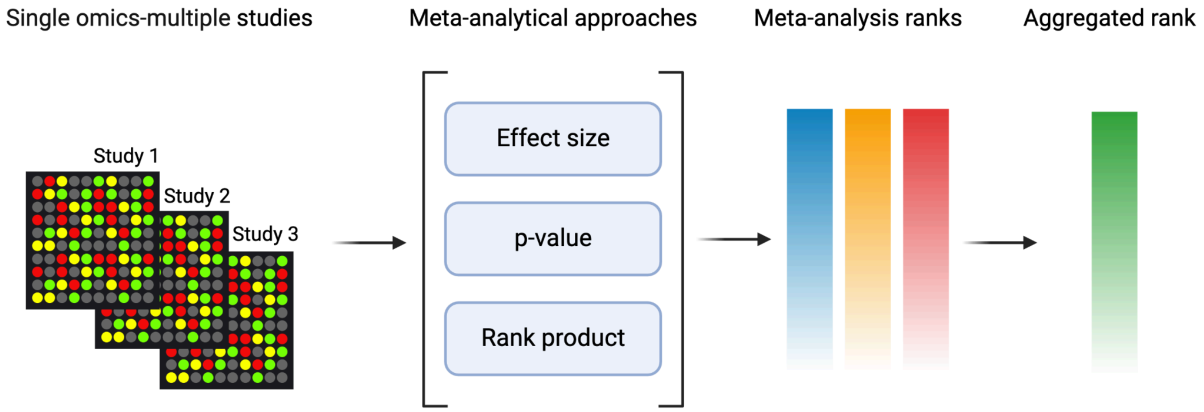


*Figure S1 - Overview of the statistical meta-analysis methods implemented in MUUMI.*

Subsequently, the user can 1) functionally characterise the gene rank obtained from the meta-analysis and 2) set a threshold on the gene rank based on biological relevance. The function *compute_gsea* is used to perform a Gene Set Enrichment Analysis (GSEA), which tests the statistical enrichment of the gene rank obtained from the meta-analysis over gene sets provided through a GMT file. The function is a wrapper of the *fgsea* function (included in the fgsea R package) and the enrichment is calculated over gene set files in GMT format that can be downloaded from the Molecular Signatures Database (MSigDB) (14).

The users should use the gmtPathways function from the fgsea package to read the GMT file specified in the input argument and create a list of gene sets. Additionally, the *plotGseaTable* function from the fgsea package (also implemented in the *compute_gsea* function) is employed to visualize the GSEA results. Eventually, the function returns the fgsea results as a data-frame containing the enrichment p-value, BH-adjusted p-value, enrichment score, normalized enrichment score, and the size of the pathway after removing genes that are not present in the gene list.

The function *compute_gsea_thresh* computes a threshold for the gene rank based on the GSEA enrichment score. The function makes use of three input parameters: geneList, which is a ranked gene list deriving from the meta-analysis; fgsea_res, which is the GSEA result obtained from the *compute_gsea* function; and background, a whole gene set object derived from the fgsea::gmtPathways(gmt_file) function. First, the function sorts the fgsea_res object by p-value in ascending order and identifies the significant pathways using a p-value cutoff of 0.05. For each significant pathway, the function calculates the GSEA enrichment score using the *calcGseaStat* function from the fgsea package. Second, the function identifies the maximum enrichment score and corresponding genes in the gene list. Finally, the function calculates the median of the maximum enrichment score indices from all the identified significant pathways and utilises it as a threshold to select the most biologically relevant features within the meta-analysis rank (Figure S2).

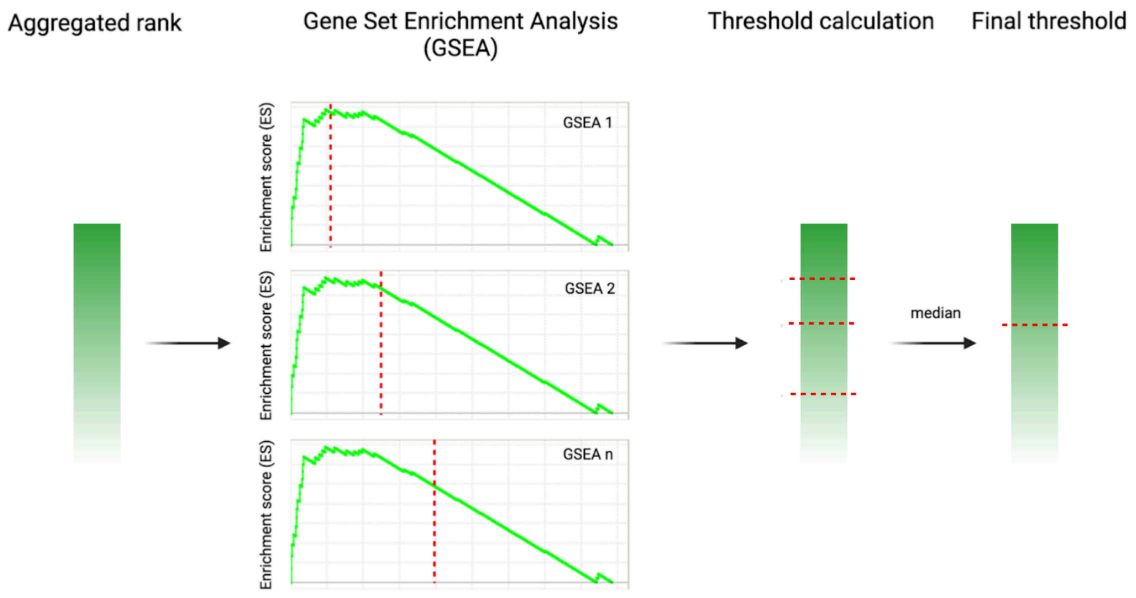


*Figure S2 - GSEA-based calculation of the meta-analysis threshold.*

**Module 2 - Inference and analysis of molecular networks.**

Molecular networks are mathematical models employed in life sciences useful to interpret the complexity of biological systems, as they represent the interactions between biomolecules such as genes, proteins, and metabolites. In addition, networks represent invaluable tools for the meta-analysis and integration of omics data, enabling researchers to extrapolate information from genomics, transcriptomics, proteomics, and metabolomics into a holistic systems-level perspective. Synthesizing molecular interactions into a structured network, facilitates the identification of key regulatory pathways, disease-associated modules, and predictive molecular targets and biomarkers, thereby supporting the interpretation of biological processes. On the other hand, integrating data deriving from different experiments poses remarkable challenges. Indeed, systematic variations in data that arise from technical differences between experiments rather than biological variability can mislead data interpretation if not taken into account. Such variation is known as “batch effect”. These effects often occur when data are collected under different experimental conditions, such as variations in sample preparation, instrumentation, or sequencing platforms. Proper identification and implementation of mitigation strategies are essential steps in data analysis, particularly in high-throughput omics studies, to ensure reliable, interpretable and reproducible results. Batch effects can result in inaccurate correlations between genes and interfere with the identification of biologically meaningful relationships (15). Therefore, batch adjustment is necessary to remove systematic variation due to technical factors, enabling a more accurate identification of true biological relationships.

For this purpose, we developed the *multi_studies_adjust* function. In this function, we embedded the *pamr.batchadjust* functionality from the pamr (PAM “Prediction Analysis of Microarrays”) package for the dataset integration. Our *multi_studies_adjust* function performs a genewise one-way ANOVA adjustment for expression values. The input to the function are omics data matrices deriving from different batches that include common genes or other molecular entities over all the batches. Assuming that sample $j$ is part of the batch $b$, and $B$ represents the entire set of samples in that batch, $x(i, j)$ represents the expression level of gene $i$ in sample $j$. The function adjusts the expression level $x(i, j)$ by subtracting the mean expression level of gene i across all samples in batch $b$, denoted by $mean[x(i, j)]$ (16). As a result, we obtain an aggregated, scaled and batch-adjusted expression matrix that can be utilised for further analyses, such as a network-based integration.

For this purpose, we developed and included in the MUUMI package a suite of functionalities that allow the inference and analysis of molecular networks. In case of single-omics multi-study datasets, the users can infer a robust, integrated network by using the output of the *multi_studies_adjust* function described before (Figure S3).

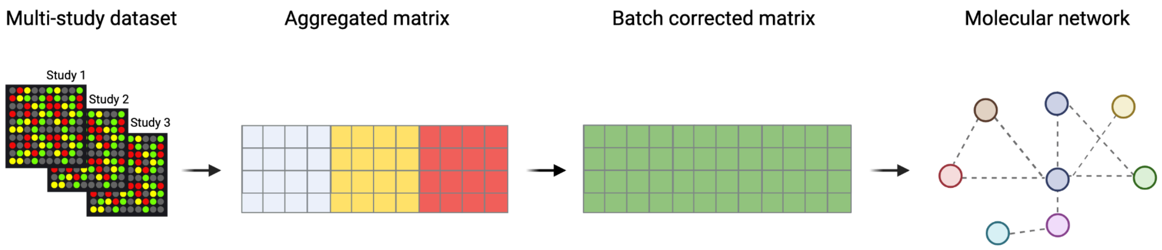


*Figure S3 - Workflow for data aggregation, batch effect mitigation and inference of molecular networks.*

To infer the network, we developed the *calculate_correlation_matrix* function, which can be used to compute the correlation between genes based on their expression levels. The function calculates the dependence between two variables (in this case, gene expression levels) by computing parametric, non-parametric and mutual information measures, starting from a gene expression table as input. Through these metrics, the function generates matrices of gene-gene correlation relationships by using all possible distinct combinations of inference algorithms, including ARACNE (17), MRNET (18), MRNETb (19) and CLR (20), mutual information, entropy estimators, and discretization methods available in the R package MINET (21), and integrated into the final network. The function allows the user to specify various parameters, such as the inference algorithms, correlation methods, and discretization methods. These parameters determine how the correlation matrix will be calculated. Once all combinations have been calculated, the function combines all the computed correlation matrices into a consensus matrix by calculating the median value for each cell in the matrices. The consensus matrix represents the overall correlation between genes, considering all the different parameter combinations. The *calculate_correlation_matrix* function is used to combine information from different methods that analyze gene expression or other omics data to create a robust molecular network deriving from an ensemble of methods. Such a network is inferred by ranking the relationships between genes based on their importance or strength. This function is based on the MINET R package (21) to create correlation matrices by mutual information methods. The user can specify the inference algorithms, correlation calculation methods and discretization methods to create multiple combinations of parameters. The outputs of multiple inferences are unified by taking their median to create a consensus matrix.

The function *get_ranked_consensus_matrix* calculates a correlation matrix for each analysis method chosen by the user in order to measure the relationships between genes (by internally calling the *calculate_correlation_matrix* function). Subsequently, the function combines the rank of edges computed through all the methods to create a single rank through the use of the *Borda* function from the TopKLists package (22). In order to complete the network inference, the user is required to run the *parse_edge_rank_matrix*. This function takes the edge rank matrix as input and selects the most important edges until all the nodes of the network are connected.

Eventually, the function gives in output a binary matrix reporting the selected edges to infer the network.
Furthermore, the user can convert the binary matrix in a more user-friendly igraph object and annotate it through the function *get_iGraph*. This function allows the user to directly annotate the graph by adding vertex attributes reporting the most important network centrality measures, including degree, betweenness, closeness, eigenvector and clustering coefficient. We also provide the users with a function to annotate a pre-made igraph object with the same centrality measures namely *annotate_iGraph*. The centrality measures can be exploited in order to get a ranked gene list through the *get_ranked_gene_list* function. This function creates gene ranks for each of the calculated centrality measures and it utilises the *Borda* function in order to compute a robust rank, where the most central genes (hubs) are included at the top.
Once the integrated graph has been created, the users can perform a community detection step through the use of the function *get_modules*. The function allows to compute a clustering on the inferred network in order to obtain biologically meaningful communities. The user can choose one of the following algorithms in order to perform the community detection: walktrap, spinglass, louvain and the fast greedy algorithm, as implemented in the igraph package.
We therefore developed functions to functionally characterise the inferred aggregated network through the analysis of the modules detected with the *get_modules* function. The package takes advantage of the Reactome database to retrieve information about biological pathways. We developed the *get_reactome_from_modules* function in order to identify significant biological pathways within each module. The function takes in input a vector containing genes that can be encoded by several gene IDs. The gene IDs that can be handled by the function can be verified through the use of the keytypes function from the org.Hs.eg.db library, and include gene symbols, Ensembl gene IDs, Ensembl Transcript IDs, Entrez IDs and Uniprot IDs, among others. The *get_reactome_from_modules* function allows the users to set the desired threshold of statistical significance by indicating the threshold of the adjusted p-value. Finally, the function writes one (or more) .csv file(s) reporting the results of the analysis. In order to visually represent the result of the functional annotation performed on the modules of the network, we developed the function *get_bubbleplot_from_pathways.* This function takes in input an igraph object reporting the modules computed on the network (possibly through the *get_modules* function) and the type of gene IDs in which the nodes of the network are encoded. The output of the function is the enrichment analysis result as a visual representation of the enriched pathways in the form of a bubble plot. The bubbles within the plot represent enriched pathways and their significance, where the bubbles are sized according to the significance level (Figure S4).

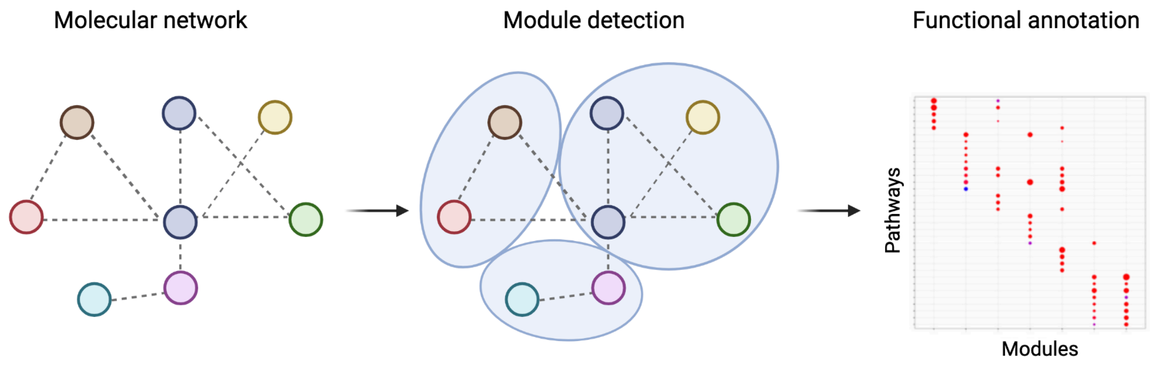


*Figure S4 - Module detection and functional analysis in molecular networks.*

While the functions for analysis and representation described until now are based on the gene content of network modules, in MUUMI we aimed to exploit topological patterns underlying network structures to uncover more precise biological interactions, as reported in the Reactome database (https://reactome.org/) or the Kyoto Encyclopedia of Genes and Genomes (KEGG; https://www.genome.jp/kegg/). These databases encompass gene-gene interactions formalising the mechanistic basis of biological processes. Here, we implemented a function aimed to perform a network-based topological functional annotation through the implementation of an Edge Set Enrichment Analysis (ESEA) function named *compute_esea*. The ESEA analysis can be performed both on the whole network or on its modules. The input to the function is a ranked vector of edges and a background of gene-gene relationships deriving from one of the following databases: KEGG (23), Reactome (24), Biocarta (25), NCI (26), SPIKE (27), HumanCyc (28) and Panther (29). For convenience, we provide the latest version of the background deriving from the KEGG database containing the reference pathway edge lists to guide the users to correctly build the background.

**Module 3 - Network integration of single-omics multi-study data**

Module 3 includes methodologies to integrate network models deriving from single-omics data deriving from different datasets. We implemented a methodology to integrate and analyze community detection results from multiple datasets, with a focus on aggregating the distinct community structures identified across heterogeneous omics data. The advantage of integrating data through the aggregation of communities is that it allows for the identification of consensus or shared patterns across datasets, enhancing the biological relevance of the results. By combining distinct community structures, the users can capture complementary information from heterogeneous omics data, reduce noise or dataset-specific biases, and uncover more comprehensive insights into the underlying molecular and functional organization of biological systems. The objective is to achieve a comprehensive integration that not only summarizes these communities but also quantifies the contributions of each dataset to the aggregated result.

This approach is particularly suited for analyzing genes within diverse omics datasets, where network communities are independently detected but require harmonization to reveal overarching patterns. To implement this strategy, we followed the framework and techniques described by Greene et al. (30), which provides a methodology for combining the results of different clustering or community detection algorithms by leveraging non-negative matrix factorization (NMF) to create unified representations and preserve dataset-specific contributions.

The *aggregate_communities* function is the entry point for this process, and it takes as input the list of communities extracted from each single dataset and the number of integrated communities to be estimated. It first constructs a stacked membership matrix representing community memberships across all datasets, using the *get_stacked_membership_matrix* function. This matrix has genes as rows and communities from all datasets as columns, with binary indicators specifying the membership of each gene in the respective community.

Once the membership matrix is prepared, the function employs a NMF model via the nmfbin package to derive an aggregated representation of communities. Specifically, it minimizes the log-loss function through gradient-based optimization, leveraging the *nndsvd* initialization to enhance convergence. The result of this step includes two outputs, the aggregated membership matrix and a set of dataset-specific contributions.

The *get_stacked_membership_matrix* function constructs the initial stacked membership matrix by iterating through community labels of each dataset. It first extracts all unique genes across datasets using the *get_unique_genes* function, ensuring consistency in gene order. It then maps the community memberships from each dataset into a binary matrix, where rows correspond to genes and columns represent communities uniquely named to reflect their dataset of origin. This ensures that all communities, even if dataset-specific, are accounted for in the aggregated analysis.

The function *compute_view_contributions* quantifies the relative contributions of each dataset to the aggregated communities derived from the NMF process. It groups the input communities by dataset origin, computes the contribution of each dataset to each aggregated community, and normalizes these contributions to account for total community weights. This step provides insights into how much each dataset informs the overall aggregation (Figure S5).

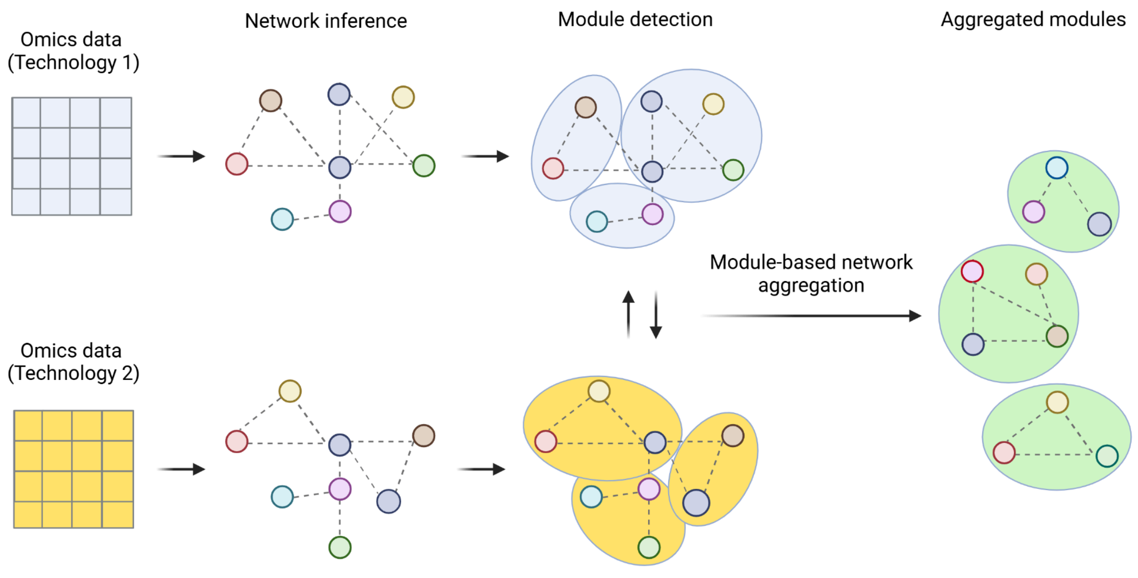


*Figure S5 - Late integration strategies based on module aggregation, as implemented in MUUMI.*

### Module 4 - Network-based integration of multi-omics data

Network-based integration of multi-omics data allows synthesizing information from diverse omics layers. This methodology organizes biomolecular entities and their interactions into heterogeneous interconnected networks, enabling the exploration of complex biological relationships and the identification of key regulatory mechanisms. By incorporating multiple data types, network-based integration captures complementary and synergistic insights that are often missed when analyzing single datasets in isolation. In MUUMI, we embedded the Similarity Network Fusion (SNF) algorithm (31), which has been designed for network-based multi-omics data integration. The aim is to combine multiple networks representing different data types (like gene expression, DNA methylation, and protein interaction data) into a unified network that reflects shared structures and relationships across the different data sources. The SNF-wrapper function included in the MUUMI package is *snf_based_integration*. Following are briefly described the integration steps of the SNF algorithm. For a more detailed explanation, please refer to (31).

1. Infer Similarity Networks

- For each data type, the algorithm infers an initial similarity network. This is done by computing the pairwise similarities between all molecular entities within each data layer.
- A common way to calculate similarity between samples is by using Gaussian similarity, which takes into account the Euclidean distance between samples while giving more weight to closer neighbors.

2. Iterative Network Fusion

- After constructing similarity networks for each data type, SNF uses an iterative process to fuse these networks. The idea is that each network will start "borrowing" information from the other networks, reinforcing connections that are consistent across data types.
- In each iteration, the similarity matrix of each network is updated by considering information from the similarity matrices of the other networks. This is often done using a process called “message passing”, which exchanges information between networks while preserving their unique structure.

3. Normalization

- Normalization typically involves re-scaling similarities to prevent one data type from dominating the fusion process.

4. Convergence to a Unified Network

- The iterative fusion continues until the networks converge, meaning that further iterations no longer change the similarity matrices significantly.
- The final result is a unified similarity network that captures the most consistent and relevant relationships between molecular entities across all data types.

The original SNF algorithm is designed to calculate similarity values among samples of a certain experiment. Our function, instead, is built to calculate similarity values among molecular entities (genes, proteins, etc.), and therefore the resulting matrix will be a squared gene-gene similarity matrix.
In such a matrix, the values represent the similarity measures among genes, forming a fully connected network where every gene is connected to every other gene. However, not all these connections are equally important or biologically meaningful. Pruning the network helps simplify it by removing weaker, less relevant connections, allowing us to focus on the strongest and most meaningful relationships. This process highlights the key interactions that are more likely to reflect true biological processes. To achieve this, we developed the *prune_snf_network* function, which retains only the most significant edges, making the network easier to interpret and more useful for downstream analysis.

19. Meyer PE, Marbach D, Roy S, Kellis M. Information-Theoretic Inference of Gene

Networks Using Backward Elimination. DBLP; 2010.
