## Supplementary material for "MUUMI: an R package for statistical and network-based meta-analysis for MUlti-omics data Integration": MUUMI case studies

### Case Study 1

#### *Experimental setup and data used*

To showcase the functionalities of MUUMI, we performed a case study on idiopathic pulmonary fibrosis (IPF) by analysing samples deriving from the integration of 17 pre-processed and harmonized public transcriptomics datasets. Among all the included datasets, 11 derive from RNA sequencing (RNA-Seq) while 6 from microarray experiments (Table S1). The collected data include both fibrotic lung biopsy samples of IPF patients and biopsy samples of healthy individuals. Datasets are made publicly available in Zenodo (https://zenodo.org/doi/10.5281/zenodo.10692128). We employed ESPERANTO, a R/Shiny application, to efficiently curate and harmonize metadata using its semi-supervised approach, streamlining the metadata curation process (1). This case study aims to perform the integration of single omics datasets and carry out both statistical and network-based meta-analysis. The resulting data can then be utilised to uncover molecular vulnerabilities and dysregulated biological processes underlying the IPF phenotype.

| GEO dataset ID | Platform | Citation | Number of IPF samples | Number of healthy samples |
|---|---|---|---|---|
| GSE150910 | Illumina NovaSeq 6000 | (2) | 103 | 103 |
| GSE213001 | Illumina HiSeq 3000 | (3) | 62 | 41 |
| GSE124685 | Ion Torrent Proton | (4) | 49 | 35 |
| GSE199949 | Illumina HiSeq 4000 | (5) | 26 | 16 |
| GSE92592 | Illumina HiSeq 2000 | (6) | 20 | 19 |

| | | | | |
|---|---|---|---|---|
| GSE199152 | Illumina HiSeq 2500 | Unpublished | 20 | 4 |
| GSE53845 | Agilent-014850 4x44K G4112F | (7) | 40 | 8 |
| GSE166036 | Illumina HiSeq 4000 | (8) | 10 | 4 |
| GSE184316 | Ion Torrent Proton | Unpublished | 40 | 24 |
| GSE110147 | Affymetrix 1.0 ST Array | (9) | 22 | 11 |
| GSE21369 | Affymetrix U133 Plus 2.0 | (10) | 11 | 6 |
| GSE24206 | Affymetrix U133 Plus 2.0 Array | (11) | 17 | 6 |
| GSE72073 | Affymetrix Transcriptome Array 2.0 | (12) | 5 | 3 |
| GSE169500 | Ion Torrent PGM | (13) | 20 | 10 |
| GSE99621 | Illumina HiSeq 2500 | (14) | 18 | 8 |
| GSE10667 | Agilent-014850 4x44K G4112F | (15) | 31 | 15 |
| GSE138283 | Illumina NextSeq 550 | (16) | 12 | 5 |

*Table S1 - Publicly available transcriptomics datasets included in the case study.*

#### *Phenotypic and molecular basis of IPF*

IPF is a chronic, progressive, devastating, and age-related interstitial lung disease (ILD) that is irreversible and typically fatal. IPF has an unknown origin and only a few treatment options that are limited in their efficacy (17). In IPF, progressive formation of scar tissue in the supportive interstitium of the lungs occurs, causing breathing to become increasingly difficult (18). Histopathologically IPF typically exhibits subpleural and paraseptal fibrosis, honeycombing, and regions of less affected and normal parenchyma (17). The proposed pathogenesis connects IPF to a lung susceptible to aging-related changes, which is subject to repetitive alveolar injuries triggered by factors such as inhaled cigarette smoke, microaspiration, nanomaterials, gastroesophageal reflux, or viruses (17,19). These injuries can lead to type I and type II epithelial cell death. Following micro-injuries and epithelial cell apoptosis, there is an increased vascular permeability to proteins like fibrinogen and fibronectin, leading to the formation of a wound clot. Subsequently, there is migration and proliferation of bronchiolar and alveolar epithelial cells, representing a rapid response of the respiratory tissue attempting self-repair. Abnormally activated epithelial cells release various chemokines, cytokines, and epidermal growth factors, attracting fibroblasts, immune system cells such as alveolar macrophages, and monocytes that differentiate into macrophages. Moreover, these cells secrete TGF-β1, promoting epithelial-mesenchymal transition, extracellular matrix remodeling, and fibroblast differentiation into myofibroblasts. The heterogeneous macrophage population also secretes chemokines and growth factors like TGF-β, inducing the growth of fibrotic tissue in the extracellular matrix. Positive feedback loops contribute to the progressive expansion of the fibrotic tissue (20–22).

#### *Identification of robust IPF genes by statistical meta-analysis*

For each dataset indicated in Table S1, we conducted differential gene expression analysis between IPF and healthy samples. For RNA-Seq datasets, gene expression data normalisation and differential gene expression analysis were performed using DESeq2 version 1.24.0 (23). DNA microarray data preprocessing was carried out by using the eUTOPIA software, which implements the limma package for expression value normalisation and performed differential gene expression analysis by applying quantile normalisation and linear model (24, 25). We adjusted nominal p-values in both RNA-Seq and microarray datasets by using the Benjamini & Hochberg method (26). In order to identify robust IPF-associated genes, we integrated the results of the differential expression analysis by performing a statistical-based meta-analysis utilising the function "*run_ensembl_metanalysis*", whose input consists of adjusted p-values from all the datasets, with gene symbols as row names. All three methods included in our pipeline, namely "effect_size," "pvalue," and "rank_product," were utilized in the ensemble analysis. 9371 genes were included in the analysis.
Among the highest-ranked genes, we identified several members of the *COL* family: *COL3A1*, which holds the highest overall ranking in the analysis, along with *COL14A1, COL15A1*, and *COL1A2* which are also in the top 20. The transcriptional profile of lung fibroblasts can influence the extracellular matrix (ECM) proteins encoded by these genes, leading to alterations in the ECM of lungs affected by IPF (27). Accumulating evidence suggests that the mechanical interactions between fibroblasts and the stiffened ECM create a feedforward mechanism, contributing to the persistence and progression of pulmonary fibrosis (28,29). A study by Huang, S. et al. 2022 (30) revealed that Asporin (ASPN) promotes the differentiation of lung myofibroblasts induced by TGF-β by facilitating Rab11-dependent recycling of TβRI. ASPN, a member of the small leucine-rich proteoglycan family known for its crucial roles in tissue injury and regeneration, is highly expressed in various tumor types and has the capacity to upregulate TGF-β1. TGF-β induces fibrotic tissue growth in the ECM (31). Elevated protein levels of MMP7 have been observed in IPF compared to samples from healthy controls, and it serves as a predictive biomarker for disease progression in IPF (32). In individuals affected by IPF, the expression of matrix metalloproteinases (MMPs) is dysregulated, resulting in substantial architectural remodeling in the lung microenvironment. MMPs have been considered potential therapeutic targets for IPF (33). Yu, G., et al. 2018 (34) found that the activity and expression of iodothyronine deiodinase 2 (DIO2), was higher in the lungs of patients with IPF compared to control individuals and correlated with disease

severity. Based on these findings, we can confirm that the genes ranked by the *run_ensembl_metanalysis* function hold biological relevance in IPF, indicating the efficacy of our approach in prioritizing genes resulting from differential analysis of multiple omics datasets.

#### *Identification of functionally relevant genes and pathways derived from statistical-based meta-analysis*

We performed Gene Set Enrichment Analysis (GSEA) utilizing the function *compute_gsea* for the ranked gene list that was obtained from the meta-analysis. Gene sets were retrieved from the Molecular Signatures Database (MSigDB) (35). To ensure meaningful results, the minimum pathway size was set to 10 genes and the number of permutations to 100,000. The 15 most enriched pathways are represented in Table S2.

| Pathway | p-val | p-adj | ES | NES | nMoreExtreme | Size |
|---|---|---|---|---|---|---|
| Extracellular Matrix Organization | 1.00E-05 | 0.003423 | 0.458414 | 1.683331 | 0 | 185 |
| Regulation of Insulin Like Growth Factor IGF Transport and Uptake by Insulin Like Growth Factor Binding Proteins IGFBPs | 1.00E-05 | 0.003423 | 0.502973 | 1.716711 | 0 | 64 |
| Collagen Degradation | 1.00E-05 | 0.003423 | 0.587627 | 1.894219 | 0 | 37 |
| Assembly of Collagen Fibrils and Other Multimeric Structures | 2.00E-05 | 0.005132 | 0.537653 | 1.763733 | 1 | 43 |
| ECM Proteoglycans | 4.00E-05 | 0.007379 | 0.512437 | 1.704437 | 3 | 49 |
| Collagen Biosynthesis and Modifying Enzymes | 5.00E-05 | 0.007379 | 0.527783 | 1.731356 | 4 | 43 |
| Molecules Associated with Elastic Fibres | 5.03E-05 | 0.007379 | 0.633992 | 1.930158 | 4 | 24 |
| Degradation of the Extracellular Matrix | 7.00E-05 | 0.00802 | 0.454145 | 1.58119 | 6 | 81 |
| Collagen Chain Trimerization | 7.03E-05 | 0.00802 | 0.603954 | 1.860309 | 6 | 26 |
| Reactome Collagen Formation | 8.00E-05 | 0.008208 | 0.482114 | 1.63859 | 7 | 61 |
| Metabolism of Steroids | 9.00E-05 | 0.008394 | 0.449728 | 1.573069 | 8 | 86 |

| | | | | | | |
|---|---|---|---|---|---|---|
| Integrin Cell Surface Interactions | 1.10E-04 | 0.009405 | 0.489075 | 1.642485 | 10 | 54 |
| Elastic Fibre Formation | 1.60E-04 | 0.012312 | 0.565026 | 1.774937 | 15 | 30 |
| Class A1 Rhodopsin Like Receptors | 1.70E-04 | 0.012312 | 0.438244 | 1.536992 | 16 | 89 |
| GPRC Ligand Binding | 1.80E-04 | 0.012312 | 0.405372 | 1.454706 | 17 | 124 |

*Table S2 – Top 15 enriched Reactome pathways in GSEA analysis.*

The *compute_gsea_thresh* function computes a threshold for the statistical meta-analysis gene rank based on the GSEA enrichment score. In this case study, the threshold was set to the top 2860 genes out of 9371 genes.
The GSEA analysis results of MUUMI (Table S2) successfully identified significant functional pathways associated with IPF. Notably, pathways related to ECM remodeling were found to be significantly enriched among the ranked genes in the meta-analysis. This emphasizes the prevalent occurrence of ECM remodeling in lung diseases such as IPF (36, 37). "Extracellular Matrix Organization" and "Collagen Formation" pathways highlight the structural aspects of the ECM, emphasizing the importance of collagen assembly and fibrillogenesis in the pathogenesis of IPF. Additionally, pathways such as "Collagen Degradation" and "Degradation of The Extracellular Matrix" imply dysregulation in the turnover of ECM components, possibly contributing to fibrotic changes observed in IPF. Pathways like "Regulation of Insulin-Like Growth Factor IGF Transport" and "Integrin Cell Surface Interactions" suggest potential connections between growth factor signaling and cellular adhesion, emphasizing the complex interplay between signaling pathways and fibrotic processes. Moreover, pathways related to elastic fibers and G protein-coupled receptor (GPCR) ligand binding shed light on the involvement of these structural and signaling elements in the context of IPF pathology. The enrichment of pathways associated with metabolism, such as "Metabolism of Steroids," suggests a broader impact on cellular homeostasis. The results of the GSEA analysis demonstrate that MUUMI is capable of recapitulating biologically relevant pathways based on the statistical meta-analysis.

#### *Mitigation of technical effects prior multi-study data integration*

Technical nuisance can heavily affect the results of omics data, leading to artifacts in the results and their interpretation. Such a phenomenon is known as batch effect. Batch effect can derive from protocols, reagents, equipment, laboratory conditions, sample collection, storage, preparation methods, time, and so forth. Therefore, since gene expression datasets utilised in our case study derive from different studies, it is essential to mitigate the batch effect before carrying out subsequent integrative analyses, such as inferring network relationships (38). To carry out batch adjustment, considering the diverse origins of the gene expression datasets, it was essential to carry out the dataset integration separately for DNA microarray and RNA-Seq datasets. The batch adjustment was carried out with the “*multi_studies_adjust*” function which takes as an input the integrated expression matrix, the sample labels, and the batch labels. The embedded functionality to carry out batch adjustment in the “*multi_studies_adjust*" utilizes the “*pamr.batchadjust*” from the "*pamr*" package (39). Figure S1 shows the outcome of the batch effect adjustment on both RNA-Sequencing (A) and DNA microarray (B) datasets. Panels A1 and B1 represent the disease samples while panels A2 and B2 represent the healthy samples.

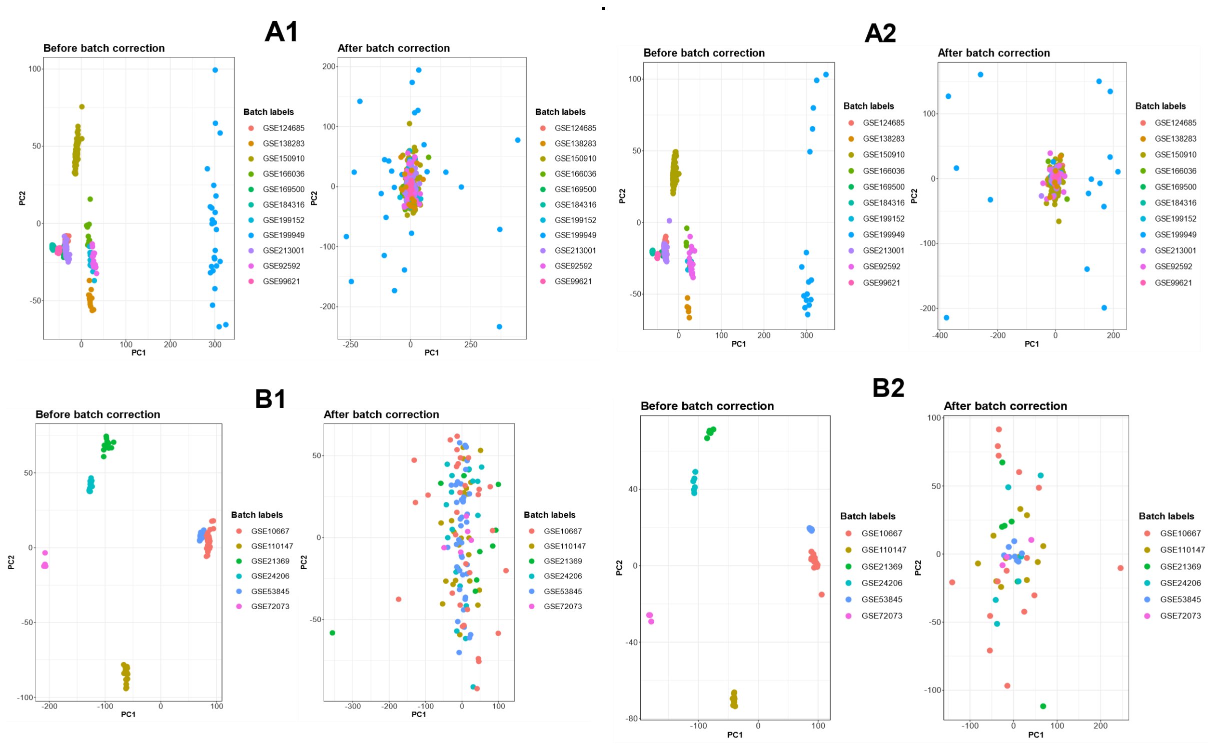


*Figure S1 - A1 and A2 PCA plots represent RNA-Seq dataset samples both before and after batch correction, with the A1 charts corresponding to disease samples and A2 charts representing healthy samples. In contrast, B1 and B2 PCA plots represent microarray dataset samples before and after batch correction, with B1 diagrams representing disease samples and B2 diagrams illustrating healthy samples.*

#### *Integrated Datasets for Network-Based Meta-Analysis*

Subsequently, we aimed at investigating co-expression patterns of the genes obtained from the meta-analysis. To do so, we inferred gene co-expression network models from IPF samples and healthy counterparts profiled through RNA-Seq datasets. To infer the networks from the normalised and batch-adjusted read count matrices, we firstly utilised the "*get_ranked_consensus_matrix*" function. Through this function we determined correlation measures between genes, based on their expression levels for both disease and healthy samples independently. In this case study, we computed Pearson correlation to build a squared adjacency matrix underlying the network and then we set CLR as a network inference algorithm. We then completed the network inference through the "*parse_edge_rank_matrix*" function to systematically add edges to the graph so that all the nodes are connected. The output of this function is a binary squared matrix, where 1 indicates the presence of an edge between two nodes while 0 indicates that no edge is present between two nodes. We therefore converted the binary matrix to an iGraph object using the *"get_iGraph"* function. The number of vertices and edges in each network are represented in Table S3.

#### *Functional characterisation of the networks*

Community detection for each network was performed using the "*get_modules*" function, which implements the walktrap (40), spinglass (41, 42), louvain (43) and the greedy algorithm (44). For this case study, we employed the walktrap algorithm. The number of communities in each network is shown in Table S3. We conducted a functional annotation for disease and healthy modules using the "*get_reactome_from_modules*" function to identify significantly enriched biological pathways. To visualize the pathway enrichment results, we used the function "*get_bubble_plot_from_pathways*" to create bubble plots for each module, with the RNA-Seq network results displayed in Figure S2. Additionally, the "*comparative_heatmap_for_modules*" function was employed to plot both healthy and disease network modules on the same plot, with the comparative bubble plots for RNA-Seq networks shown in Figure S3.

| **RNA-seq disease network** | | | **Microarray disease network** | | | **RNA-seq healthy network** | | | **Microarray healthy network** | | |
|---|---|---|---|---|---|---|---|---|---|---|---|
| **Modules** | **Vertices** | **Edges** | **Modules** | **Vertices** | **Edges** | **Modules** | **Vertices** | **Edges** | **Modules** | **Vertices** | **Edges** |
| 6 | 10460 | 899649 | 17 | 14437 | 612359 | 8 | 10460 | 1373581 | 6 | 14437 | 572961 |

*Table S3 – Number of modules, vertices, and edges in RNA-Seq and microarray networks.*

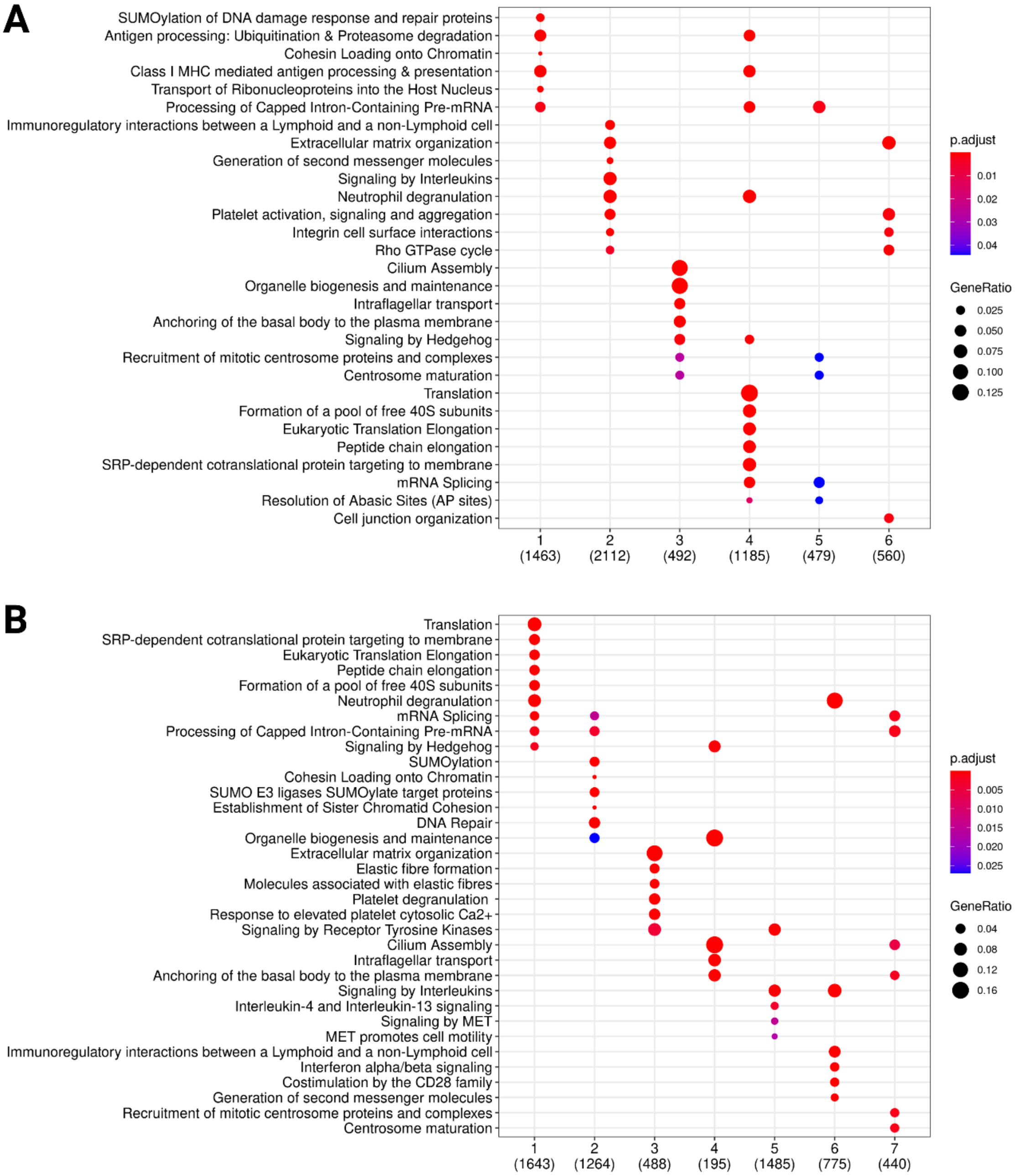


*Figure S2 – Bubble plots of significant Reactome pathways in the RNA-Seq disease network (A) and RNA-Seq healthy network (B). X-axis represents the communities in the networks and Y-axis represents the Reactome pathways. The color of the bubble represents the significance of the pathway in the module and the size of the bubble represents the relative amount of genes in the module that are present in the Reactome pathway.*

The network derived from the meta-analysis of RNA-sequencing data identified several modules associated with disease pathology. One of the primary disease-associated modules involves antigen presentation and processing, emphasizing the role of immune system activation in fibrosis (45). Also, immunoregulatory interactions between a Lymphoid and a non-Lymphoid cell Reactome pathway were significantly enriched in disease module two. Different chemokines are connected to the occurrence of IPF, and an uneven distribution of angiogenic chemokines has been correlated with vascular remodeling in the condition (45). Another significant module includes platelet activation, neutrophil degranulation, and extracellular compartment alterations, suggesting a strong inflammatory response and extracellular matrix remodeling, both critical in disease development. The ECM in the lungs consists of collagens, elastin, glycoproteins, and proteoglycans. These components play a central role in providing structural support for cells and ensuring the mechanical stability and elastic recoil essential for normal lung function. In IPF, a characteristic feature is the accumulation of myofibroblasts in clusters known as fibroblastic foci. This leads to a significant deposition of ECM within the interstitium, destroying lung architecture (46). Additionally, cilia impairment has been identified as a key factor, reinforcing its relevance in various respiratory conditions. The analysis also highlights significant alterations in translation, transcription, and mRNA metabolism, indicating broad regulatory changes at the molecular level that could contribute to disease-associated gene expression alterations. Furthermore, module five underscores disruptions in cell junction integrity and its links to the extracellular functions reported in module 2.

In contrast, the modules identified in healthy conditions highlight key biological processes crucial for homeostasis, including translation, transcription, and mRNA metabolism. Elastic fibers also emerge as an essential module, emphasizing their role in tissue elasticity and structural integrity. Notably, the control transcriptional profiles derive from healthy lungs, which therefore retain the majority of tissue functionalities. Cilia function remains a key feature in healthy conditions, mirroring its significance in disease when this function is altered.

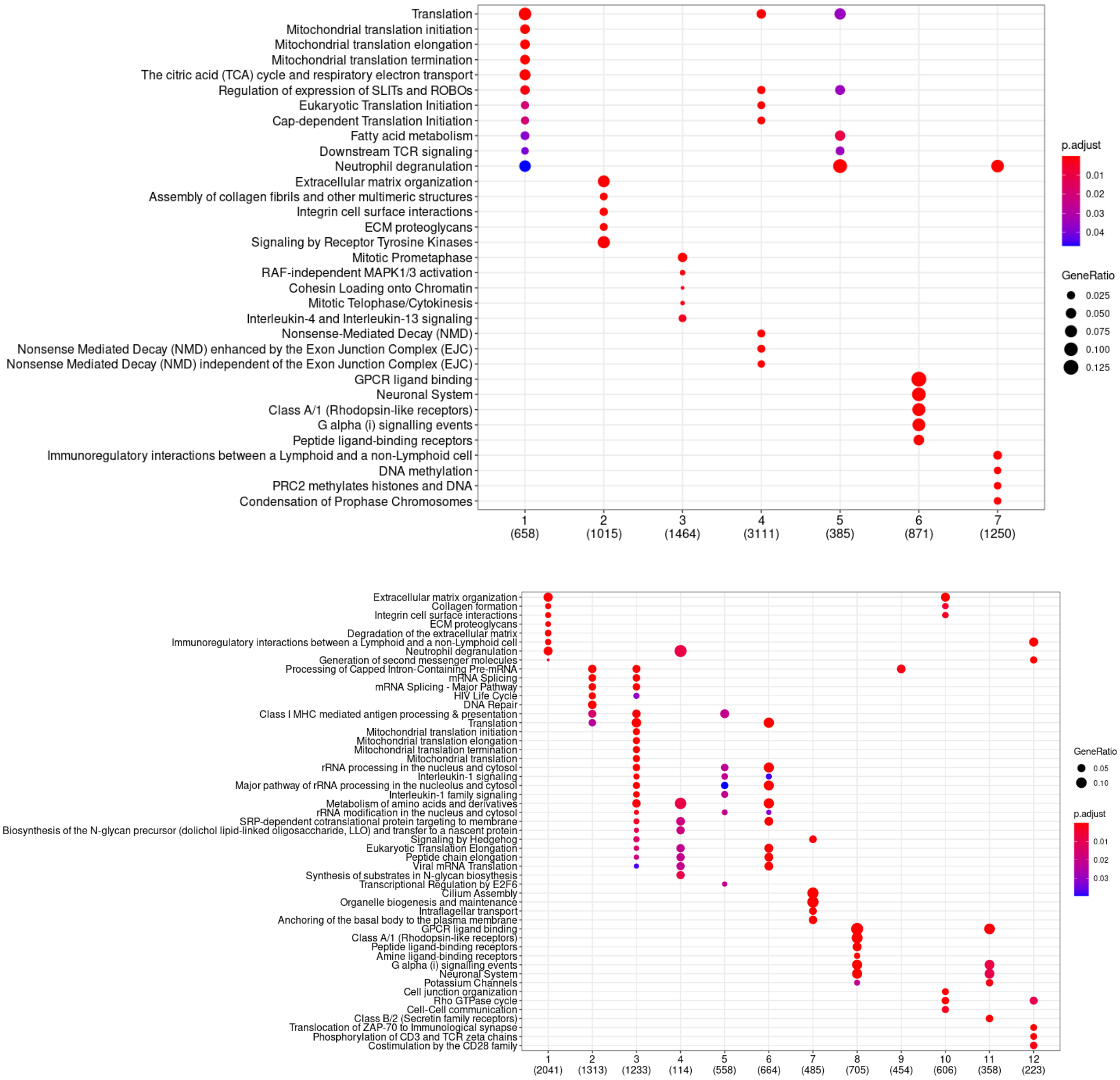


*Figure S3 – Bubble plots of significant Reactome pathways in the integrated co-expression networks deriving from both RNA-Seq and microarrays. Panel A refers to the communities deriving from IPF samples and panel B refers to communities deriving from healthy samples. X-axis represents the community numbers and Y-axis represents the enriched pathways. The color of the bubble represents the significance of the pathway in the module and the size of the bubble represents the relative amount of genes in the module that are present in the pathway.*

Subsequently, we showcase the late integration functionalities of MUUMI by integrating transcriptomics data deriving from both RNA-Seq and DNA microarrays. First, we identified the communities in each of the inferred networks for both of the technologies. We then used the "*aggregate_communities*" function to independently aggregate communities from RNA-Seq and DNA microarray experiments for both disease and healthy networks. The number of modules for the aggregated communities was determined by averaging and rounding to the nearest integer based on the module counts in the RNA-Seq and DNA microarray networks Table S3.

By incorporating DNA microarray data alongside RNA-Seq, the merged meta-analysis yielded cleaner and more consolidated results. Notably, functional redundancies observed in RNA-Seq disease modules, such as modules 2 and 5, appear condensed into a single module, streamlining the interpretation of immune-related and extracellular processes. Previously unobserved functions, such as the impact on mitochondria, have now emerged as key elements. This finding aligns with recent research demonstrating that mitochondrial quality significantly influences pulmonary fibrosis (PF) progression (47). Further considerations suggest that mitochondrial function plays a crucial role in disease progression. The emergence of mitochondrial dysfunction as a significant factor provides a novel perspective on pulmonary fibrosis pathogenesis. This insight is supported by recent findings highlighting mitochondrial quality as a key determinant of disease severity. Additionally, interleukin signaling, particularly IL-4 and IL-13, plays a crucial role in immune regulation and tissue maintenance. Furthermore, interferons, CD28, and other interleukins emerge as critical immune modulators maintaining a healthy physiological state.

Additionally, the role of nonsense-mediated mRNA decay (NMD) has been identified in module 4, implicating various mechanisms of gene expression regulation. NMD serves as a critical surveillance pathway for controlling mRNA quality and abundance. Gene expression deregulation through various epigenetic mechanisms is a known contributor to idiopathic pulmonary fibrosis (IPF). Interestingly, NMD based approaches have recently emerged as a therapeutic target in cystic fibrosis.

This functional interpretation offers valuable insights into the biological processes underlying disease states. The integration of multiple datasets not only refines the understanding of previously identified functionalities but also reveals novel functional associations, particularly with mitochondrial dysfunction and RNA surveillance mechanisms. Overall, the diverse array of pathways highlights the intricate molecular landscape involved in IPF, providing potential targets for therapeutic intervention and a deeper understanding of the disease mechanisms.

Finally, we computed the contributions of each of RNA-Seq and microarray technologies to the inference of the integrated communities. The contribution of each data source is calculated as the sum of its weights across all genes, normalized by the total weight for each community. These weights are derived using non-negative matrix factorization (NMF), which integrates gene communities identified in each data source and condition. Notably, some integrated communities exhibit relatively balanced contributions from all data sources (e.g., communities 1–7 in the diseased samples and community 1 in the healthy samples), indicating that these communities are shared across multiple data views. In contrast, the remaining communities show more uneven contributions, suggesting that these integrated communities are uniquely driven by a single data source. These findings underscore the strength of this integrative approach in both consolidating redundant communities across different data modalities and capturing unique, dataset-specific information that would otherwise remain undetected.

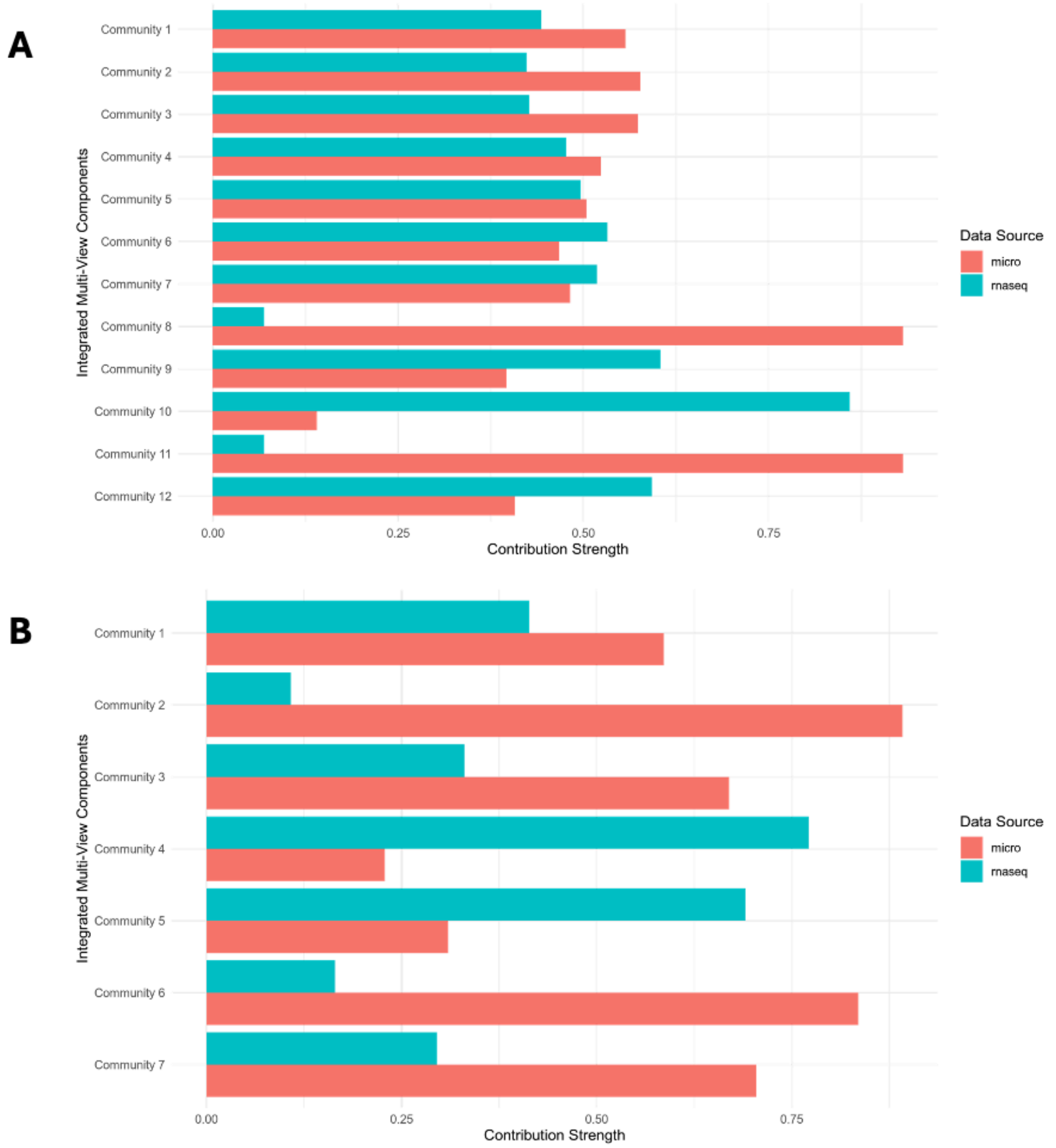


*Figure S4 - Contribution of each individual data source to the integrated communities in both disease (panel A) and healthy (panel B) samples.*

### Case Study 2

#### *Multi-omics data integration to uncover the plastic nature of macrophages upon polarization stimuli*

This case study demonstrates the potential of MUUMI, to integrate multi-omics datasets. Integrating multi-omics data is essential to gain a comprehensive understanding of complex biological systems, as it reveals the intricate regulatory mechanisms operating across different molecular layers. Combining transcriptomics, epigenomics, proteomics, and other omics layers provides insights into biological system regulation that single-omics approaches may overlook (48).

In this case study, we analyzed publicly available bulk RNA-Seq and DNA methylation array data previously collected from Gene Expression Omnibus (GEO), reported with the GEOID: GSE273628. In this study, Migliaccio et al. (49) explored the regulatory relationship between DNA methylation and gene expression in macrophages exposed to different polarization stimuli. Cells were differentiated with 30.9 ng/ml of phorbol 12-myristate 13-acetate (PMA) for 48h. After PMA differentiation, cells are exposed to two different cytokines cocktails and fresh media for 24h, 48h, and 72h. In particular, to simulate a pro-inflammatory environment the cells are exposed to LPS 10 pg/ml and INFγ 20 ng/ml; to induce an anti-inflammatory phenotype instead, the cells are exposed to IL-13 20 ng/ml and IL-4 20 ng/ml.

To gain a comprehensive understanding of the capabilities of the tool and its potential to uncover mechanisms underlying molecular responses to different polarization stimuli, we analyzed the epigenomic and transcriptomic profiles of macrophages stimulated with LPS-INFγ and IL4-IL13. To comply with the findings of Migliaccio et al. 2024, which emphasize the slower kinetics of DNA methylation compared to transcriptional changes, we selected the 72-hour time point for our analysis. This timeframe allows for the detection of more established epigenetic modifications, including those that may have originated at earlier time points.

To integrate these datasets, we utilized the *snf_based_integration* function implemented in MUUMI. Through our analysis, we generated two fusion networks: one for the LPS-INFγ and one for the IL4-IL13-treated macrophages. Such networks are completely connected, and each edge indicates the similarity values between genes. The similarity parameter assesses the degree of concordance between the expression patterns of gene pairs in both the transcriptomic and epigenomic layers. A higher similarity score suggests that the genes in a pair tend to exhibit similar regulatory patterns in transcriptomics and epigenomics layers. Both of the networks are composed of 12,940 nodes and 163,443,600 edges. To retain only the most biologically meaningful connections and make the computation faster, we pruned both of the networks through the function *prune_snf_network,* by setting the 90$^{th}$ percentile as a threshold of significance. The pruned LPS-INFγ network encompassess 8,502,486 edges while the IL4-IL13 one 8,500,640 edges.

Subsequently, we ranked the network edges based on their weights and performed a functional annotation using KEGG pathways through ESEA (Enrichment Set Enrichment Analysis). By performing an enrichment analysis on edges with high similarity scores, we can uncover biological processes that are strongly coordinated at gene expression and epigenetic level. To identify the most significant biological processes and pathways associated with each condition, we performed separate enrichment analysis on the LPS-INFγ and IL4-IL13 networks. The most significant enriched terms (adjusted pvalue <0.05) are presented in Figure S5.

Our integrated network analysis of the LPS-INFγ condition reveals a strong correlation between genes involved in TLR-receptor and TNF signaling pathways (50), consistent with the findings of Migliaccio et al. at the transcriptome level. LPS activates TLR4, initiating MyD88-dependent signaling that drives NF-κB activation and TNF-α expression, amplifying inflammation through autocrine and paracrine effects (51).

In addition, this integrated network further demonstrates the crucial role of metabolic pathways in gene regulation (52, 53), as also observed by Migliaccio et al. Notably, genes implicated in chronic inflammatory diseases, such as rheumatoid arthritis, also exhibit high similarity within this integrated network. The presence of these epigenetic modifications suggests that alterations in the epigenetic regulation of

these genes may contribute to the development and maintenance of chronic inflammatory states. In contrast to the LPS-INFγ network, fewer pathways appear enriched within the IL4-IL13 network, demonstrating a milder response to IL4-IL13 stimulation compared to LPS-INFγ. Nevertheless, the presence of terms related to immune pathways suggests the existence of distinct gene groups exhibiting high correlation across both regulatory layers within this integrated network.
By facilitating network-based meta-analyses, MUUMI offers a versatile platform that enables researchers to identify crucial biological pathways, uncover synergistic regulatory effects, and effectively pinpoint key molecular targets. By employing the network analysis features of MUUMI, researchers can integrate omics networks, providing valuable insights into gene sets exhibiting high concordance in all regulatory layers.

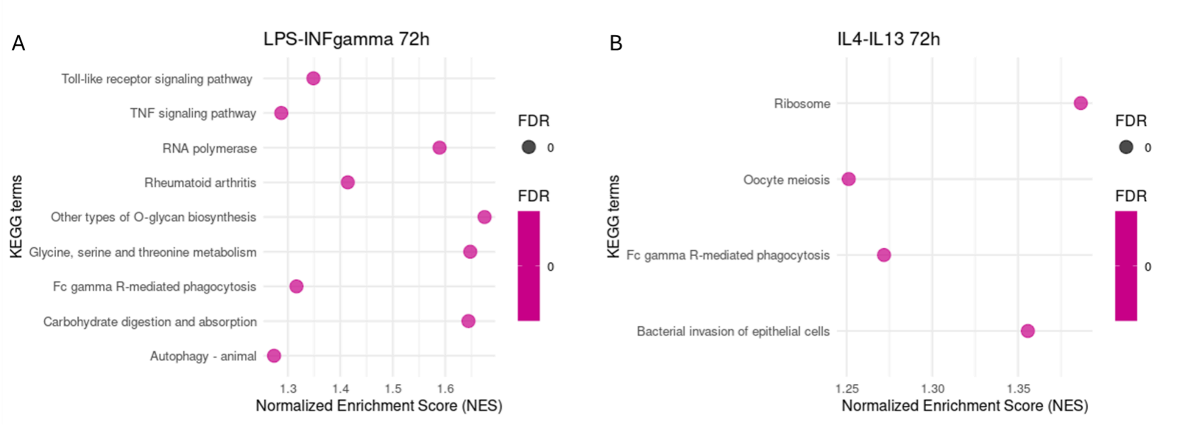


*Figure S5 - Results of the Edge Set Enrichment Analysis (ESEA) on the multi-omics integrated networks of LPS-INFγ-treated macrophages (panel A) and the IL4-IL13-treated macrophages (panel B).*

#### *Comparison of MUUMI with existing tools*

In this study, we compared the functionalities of MUUMI with other similar packages or web services dedicated to meta-analysis and integration of omics data. We found that, to date, there are 5 tools that belong to the same thematic area of MUUMI.
Tools that integrate statistical meta-analysis with network-based meta-analysis for diverse omics data remain scarce. Existing tools often lack the flexibility for users to customize analytical pipelines to suit specific research questions. This limitation stems from restricted code accessibility, rigid pipelines, and inflexible file formats. MUUMI proposes a toolbox that integrates 1) a robust ensemble-based meta-analytical approach, 2) a novel data-driven feature selection function to prioritise biologically meaningful features, 3) tools to manipulate and interpret molecular networks. These functionalities can be applied both to single and multi-omics data deriving from multiple studies. MUUMI's R functions are made publicly available and the users can build a completely tailored pipeline, by adapting it to the specific biological problem under investigation.
Among the investigated tools, MetaOmics, developed by (54), is the most similar software to MUUMI in terms of functionalities but , to date, it is designed only for transcriptomics data, including bulk RNA-Seq and DNA microarray data. MetaOmics provides a comprehensive suite for meta-analyzing transcriptomic studies, encompassing metadata quality control, differential expression analysis, pathway enrichment, differential co-expression network analysis, and network modules integration. In detail, MetaOmics offers a web interface built in R Shiny. While that makes it user-friendly for non-coders, it restricts customization beyond the provided graphical interface. The suite includes several specialized modules, each targeting a specific aspect of transcriptomic meta-analysis. For instance, MetaQC provides quantitative measures to assess the quality of individual studies, supporting the user in dataset inclusion or exclusion for meta-analysis. MetaDE implements various statistical methods to identify differentially expressed genes across multiple studies, enhancing the detection of consistent gene expression changes. MetaPath focuses on functional annotation by integrating results from multiple studies, giving insights into the biological processes associated with the biological conditions under investigation. MetaNetwork analyzes co-expression networks to identify changes in gene interactions between different conditions, offering a network-level understanding of transcriptomic alterations. However, the current implementation lacks the ability to rank genes based on centrality measures and perform analyses that leverage the topological features of the network. MetaPredict develops predictive models by combining data from multiple studies in order to improve the robustness of predictions related to disease outcomes or other phenotypes. MetaClust performs clustering analyses for disease subtype identification or gene clusters by integrating data across studies. Finally, MetaPCA applies principal component analysis in a meta-analytic framework to reduce data dimensionality, providing also an instrument for visualization and interpretation of complex transcriptomic data.
MixOmics (55) is a comprehensive R package designed for the integration and multivariate analysis of high-dimensional omics data, such as genomics, transcriptomics, proteomics, and metabolomics. It provides a suite of statistical methods to explore relationships between multiple datasets, enabling researchers to identify biomarkers, detect latent structures, and uncover biological insights from

complex data. MixOmics supports various unsupervised and supervised approaches, including Principal Component Analysis (PCA), Partial Least Squares (PLS), Canonical Correlation Analysis (CCA), and DIABLO (Data Integration Analysis for Biomarker discovery using Latent variable approaches for Omics studies). The package offers tools for feature selection, classification, clustering, and network inference, facilitating a holistic understanding of biological systems. Additionally, MixOmics includes powerful visualization functions, such as correlation networks and sample plots, to help interpret results intuitively. MixOmics is among the most similar tools to MUUMI, allowing the meta-analysis and integration of multi-omics datasets through a number of statistical tools, along with network inference. Like in the case of Netmeta, Mixomics does not include any tool for the exploitation of network topology for biological interpretation.

Mergeomics (56) is a Bioconductor package that allows the integration of multidimensional biological data. It requires an existing network as input but it does not offer functionalities for network inference. Key functionalities of Mergeomics include Marker Dependency Filtering (MDF), which corrects for known dependencies between omics markers, such as linkage disequilibrium between single nucleotide polymorphisms (SNPs), ensuring that redundant information does not bias downstream analyses. Marker Set Enrichment Analysis (MSEA) identifies disease-relevant biological processes by integrating omics markers with functional genomics data, canonical pathways, or co-expression networks, helping detect biological processes perturbed in disease by integrating multi-omics and prior knowledge. Key Driver Analysis (KDA) pinpoints essential regulators of disease-associated pathways and networks by analyzing the topology of biological networks, identifying local hubs whose neighbors are enriched for genes in the disease-associated gene sets, thereby highlighting potential key regulators. Moreover, Mergeomics includes a module able to predict therapeutic drugs by matching multi-omics informed disease pathways or networks with drug signatures. This functionality is particularly interesting for researchers working in the systems pharmacology field and are seeking for drug discovery or drug repurposing predictions.

The Netmeta package (57) is a R tool designed for network meta-analysis, which extends traditional meta-analysis by comparing multiple treatments simultaneously across a network of studies. It enables researchers to synthesize direct and indirect evidence, providing estimates of relative treatment effects even when some comparisons have not been studied head-to-head. Netmeta supports frequentist network meta-analysis, offering functionalities such as automated construction of network graphs, estimation of treatment rankings, assessment of inconsistency between direct and indirect evidence, and sensitivity analyses. While Netmeta includes several functions dedicated to network meta-analysis, its application does not lie in the domain of omics data analysis or integration. Netmeta does not include functionalities for centrality-based functional annotation or Edge Set Enrichment Analysis, limiting the possibility for biological interpretation of network structures.

MOFA (Multi-Omics Factor Analysis) (58) is a statistical framework designed for the integration and dimensionality reduction of multi-omics data in an unsupervised fashion. The tool allows to identify hidden patterns in complex datasets and to identify key factors that explain differences between samples. By implementing Bayesian factor analysis, MOFA simplifies large datasets, making it easier to uncover important biological signals, such as patient subgroups or key molecular drivers of biological processes, including diseases. The tool also supports missing data handling, allowing for incomplete datasets to be analyzed without imputation. Moreover, MOFA provides visualization functions to perform downstream association analyses, and interpret multi-omics relationships. MOFA is one of the most used tools in biomedical research, particularly for identifying disease subtypes, characterizing molecular heterogeneity, and gaining mechanistic insights from large-scale omics data analyses. It has been recently updated with a new version, named MOFA+ (59). Compared with MUUMI, MOFA and MOFA+ do not include functions for the inference, analysis and interpretation of molecular networks, neglecting the importance of the relationships among the molecular entities under investigation.

Compared to the aforementioned tools, MUUMI balances ease of usage, flexibility of the pipelines and integration of several algorithms and methods that offer a plethora of functionalities to the research community. In fact, MUUMI integrates both statistical meta-analysis with network-based methods for integration of multi-omics data. On one hand, MUUMI combines statistical methods that have been widely used, not only in the omics field, with new ways of selecting biologically relevant features. On the other hand, our package implements network-based tools to integrate single and multi-omics data in order to facilitate the biological interpretation of high-dimensional datasets. Notably, MUUMI not only integrates analyses that are scattered across different packages and web tools, but it gives a lot of emphasis on the network integration and analysis. In addition, it is the only tool among those compared that enables Edge Set Enrichment Analysis (ESEA). This is crucial because it leverages the biological significance of the interactions among the molecular entities under study, providing a remarkable resolution for interpretation of the network models. Furthermore, many available R packages address specific aspects of multi-omics data integration. However, MUUMI uniquely combines these functionalities into a single, comprehensive package giving emphasis on both data integration and network characterization. Table S5 summarizes the comparison of tools, highlighting the presence or absence of key features for each tool.

| | **MetaOmics** | **mixomics** | **MergeOmics** | **Netmeta** | **MOFA** | **MUUMI** |
|---|---|---|---|---|---|---|
| | **R shiny** | **R package** | **R package** | **R package** | **R package** | **R package** |
| **Preprocessing of transcriptomics data** | X | | | | | |
| **Metadata quality control** | X | | | | | |
| **Differential gene expression analysis** | X | | | | | |
| **Batch adjustment** | | X | | | X | X |
| **Dimensionality reduction** | X | X | | | X | X |
| **Gene-level meta-analysis** | X | | | | X | X |
| **Single omics integration** | X | X | X | X | X | X |
| **Multi-omics integration** | | X | X | X | X | X |
| **Ranking based on differential analysis** | | X | | | | X |

| | | | | | | |
|---|---|---|---|---|---|---|
| **Network centrality-based ranking** | | X | X | X | | X |
| **Network inference** | | X | X | X | | X |
| **Node functional annotation** | X | | X | X | | X |
| **GSEA or similar** | | | X | | | X |
| **ESEA** | | | | | | X |
| **SNF** | | | | | | X |
| **SNF to igraph** | | | | | | X |
| **Module aggregation of multiple network modules** | X | | | | | X |
| **Functional annotation of network modules** | X | | X | | | X |
| **Network comparison at module level** | | | | | | X |

*Table S5- Comparison of MUUMI to other similar tools.*

#### *Limitations of the study*

This study utilised multi-omics profiles obtained from public repositories. The availability of clinical data was usually insufficient. The limited detailed clinical information in many datasets manifests challenges to the predictive capacity of this study. It inhibits our ability to infer certain clinical parameters for patients with IPF, such as disease severity, time elapsed since the initial diagnosis, and the anatomical location of the samples. Additionally, the lack of information concerning the current administration of active pharmacological therapy to the patients is a noteworthy limitation. The transcriptional signatures shaping the topology of network models may be impacted by ongoing or terminated pharmacological therapies, potentially interfering with the pharmacological footprint investigated in this study. Consequently, the lack of comprehensive sample characterization restricts the translational potential of this research. The lack of randomization and data FAIRness in many datasets was problematic. The minimum standards for omics data often result in poor usability due to incomplete characterization of the experimental design and execution, as well as the lack of description regarding potential systematic effects caused by reagents, microarrays, and other factors (60).

39. T. Hastie, R. Tibshirani, Balasubramanian Narasimhan and Gil Chu. Pam: Prediction Analysis for Microarrays. R package version 1.56.1.

https://CRAN.R-project.org/package=pamr. CRAN R-project; 2019.

40. Pons P, Latapy M. Computing communities in large networks using random walks. J Graph Algorithms Appl. 2006;10(2):191–218.

41. Reichardt J, Bornholdt S. Statistical mechanics of community detection. Phys Rev E. 2006 Jul 18;74(1).

42. Newman MEJ, Girvan M. Finding and evaluating community structure in networks. Phys Rev E Stat Nonlin Soft Matter Phys. 2004 Feb 26;69(2 Pt 2):026113.

56. Ding J, Blencowe M, Nghiem T, Ha S-M, Chen Y-W, Li G, et al. Mergeomics 2.0: a web

server for multi-omics data integration to elucidate disease networks and predict therapeutics. Nucleic Acids Res. 2021 Jul 2;49(W1):W375–87.

57. Balduzzi S, Rücker G, Nikolakopoulou A, Papakonstantinou T, Salanti G, Efthimiou O, et al. netmeta : An *R* Package for Network Meta-Analysis Using Frequentist Methods. J Stat Softw. 2023;106(2).

58. Argelaguet R, Velten B, Arnol D, Dietrich S, Zenz T, Marioni JC, et al. Multi-Omics Factor Analysis-a framework for unsupervised integration of multi-omics data sets. Mol Syst Biol. 2018 Jun 20;14(6):e8124.

59. Argelaguet R, Arnol D, Bredikhin D, Deloro Y, Velten B, Marioni JC, et al. MOFA+: a statistical framework for comprehensive integration of multi-modal single-cell data. Genome Biol. 2020 May 11;21(1):111.

60. Saarimäki LA, Melagraki G, Afantitis A, Lynch I, Greco D. Prospects and challenges for FAIR toxicogenomics data. Nat Nanotechnol. 2022 Jan;17(1):17–8.
